# Identification of biomarkers for COVID-19 associated secondary hemophagocytic lymphohistiocytosis

**DOI:** 10.1101/2024.08.13.607855

**Authors:** Susan P. Canny, Ian B. Stanaway, Sarah E. Holton, Mallorie Mitchem, Allison R. O’Rourke, Stephan Pribitzer, Sarah K. Baxter, Mark M. Wurfel, Uma Malhotra, Jane H. Buckner, Pavan K. Bhatraju, Eric D. Morrell, Cate Speake, Carmen Mikacenic, Jessica A. Hamerman

**Affiliations:** Center for Fundamental Immunology, Benaroya Research Institute, Seattle, WA; Department of Pediatrics, University of Washington, Seattle, WA; Kidney Research Institute and Division of Nephrology, Department of Medicine, University of Washington, Seattle, WA; Division of Pulmonary, Critical Care and Sleep Medicine, Department of Medicine, University of Washington, Seattle, WA; Center for Translational Immunology, Benaroya Research Institute, Seattle, WA; Center for Systems Immunology, Benaroya Research Institute, Seattle, WA; Sonoma Biotherapeutics, Seattle, WA; Department of Infectious Disease, Virginia Mason Medical Center, Seattle, WA; Department of Medicine, Section of Infectious Diseases, University of Washington, Seattle, WA; Department of Immunology, University of Washington, Seattle, WA; Center for Interventional Immunology, Benaroya Research Institute, Seattle, WA

## Abstract

**OBJECTIVES:** We aimed to define and validate novel biomarkers that could identify individuals with COVID-19 associated secondary hemophagocytic lymphohistiocytosis (sHLH) and to test whether fatalities due to COVID-19 in the presence of sHLH were associated with specific defects in the immune system.

**DESIGN:** In two cohorts of adult patients presenting with COVID-19 in 2020 and 2021, clinical lab values and serum proteomics were assessed. Subjects identified as having sHLH were compared to those with COVID-19 without sHLH. Eight deceased patients defined as COVID-sHLH underwent genomic sequencing in order to identify variants in immune-related genes.

**SETTING:** Two tertiary care hospitals in Seattle, Washington (Virginia Mason Medical Center and Harborview Medical Center).

**PATIENTS:** 186 patients with COVID-19

**INTERVENTIONS:** None

**MEASUREMENTS AND MAIN RESULTS:** Nine percent of enrolled COVID-19 subjects met our defined criteria for sHLH. Using broad serum proteomic approaches (O-link and SomaScan), we identified three biomarkers for COVID-19 associated sHLH (soluble PD-L1, TNF-R1, and IL-18BP), supporting a role for proteins previously associated with other forms of sHLH (IL-18BP and sTNF-R1). We also identified novel biomarkers and pathways of COVID-sHLH, including sPD-L1 and the syntaxin pathway. We detected variants in several genes involved in immune responses in individuals with COVID-sHLH, including in *DOCK8* and in *TMPRSS15*, suggesting that genetic alterations in immune-related genes may contribute to hyperinflammation and fatal outcomes in COVID-19.

**CONCLUSIONS:** Biomarkers of COVID-19 associated sHLH, such as soluble PD-L1, and pathways, such as the syntaxin pathway, and variants in immune genes in these individuals, suggest critical roles for the immune response in driving sHLH in the context of COVID-19.

**Key Points:** *QUESTION:* To define biomarkers that could identify individuals with COVID-19 associated secondary hemophagocytic lymphohistiocytosis (sHLH) and to test whether fatalities due to COVID-19 in the presence of sHLH were associated with specific defects in the immune system.

*FINDINGS:* In two independent cohorts using two different platforms, we identified sPD-L1, IL-18BP, and sTNF-R1 as COVID-sHLH biomarkers. We identified the syntaxin pathway as important in COVID-sHLH and variants in immune-related genes in a subset of deceased COVID-sHLH subjects.

*MEANING:* Immune related proteins and pathways are dysregulated in COVID-sHLH.

## Introduction

COVID-19, caused by infection with the novel severe acute respiratory syndrome coronarvirus-2 (SARS-CoV-2), has led to over 6 million deaths worldwide. COVID-19 can be complicated by severe respiratory symptoms leading to hospitalization or acute respiratory distress syndrome (ARDS). There is increasing evidence for the role of a dysregulated immune response accompanied by elevated cytokine levels contributing to severe disease (1–5). Data suggest that some patients with severe illness present with features of a cytokine storm syndrome (CSS) (4). The term CSS encompasses several related hyperinflammatory disorders including cases of secondary hemophagocytic lymphohistiocytosis (sHLH), which can complicate infections, such as Epstein Barr Virus (EBV) or influenza. sHLH can also occur secondary to malignancies or to rheumatic diseases, where it is often referred to as macrophage activation syndrome (MAS) (1, 3). Common sHLH features include cytopenias, hyperferritinemia, coagulopathy, and end organ dysfunction. Important roles for T and natural killer (NK) cells and macrophages are well recognized in driving the pathophysiology of sHLH. A positive feedback loop is established in which interferon gamma (IFNγ) secretion by CD8+ T and NK cells activates macrophages, which secrete inflammatory cytokines, such as IL-1, IL-6, IL-12 and IL-18, to drive further lymphocyte activation (1, 6). Primary HLH (pHLH) is due to genetic defects in the T and NK cell cytolytic pathway, whereas sHLH is thought to be caused by a combination of genetic factors, induced defects in the cytolytic pathway, and inflammasome hyperactivation.

The type I interferon (IFN) cytokine family (e.g. IFNα, IFNβ) has specifically been implicated as important in COVID-19 (7–9). However, the role of IFNγ in SARS-CoV-2 infection is less clear as higher IFNγ levels have been reported in deceased subjects compared to those that survived but also in patients with moderate disease compared to severe disease and in patients with asymptomatic compared to symptomatic infections (10–12). Differences between studies may be related to the specific timing of IFNγ measurement during the course of disease as well as the accompanying inflammatory environment. For example, TNF in combination with IFNγ may provide a synergistic effect leading to hyperinflammation in models of SARS-CoV-2 infection (13). However, PD-L1, which is induced by IFNγ (14), is a potential biomarker of poor outcomes in COVID-19. Patients with higher levels of soluble PD-L1 (sPD-L1) were more likely to die from COVID-19, and sPD-L1 was also correlated with need for invasive mechanical ventilation (7, 15, 16). Together, these data suggest a role for IFNγ in severe COVID-19.

Here we generate a definition of COVID-sHLH that we used to identify individuals from available clinical data. We identified putative biomarkers of COVID-sHLH and proteins and pathways associated with COVID-sHLH. Genetic analyses in fatal COVID-19 suggest that defects in immune pathways may contribute to severe or fatal outcomes, although we did not find any variants in HLH-associated genes in our subjects.

## Methods

### Study population and design

We conducted a prospective cohort study of SARS-CoV2 positive patients enrolled at two hospital systems in Seattle, WA in 2020 and 2021. Samples for cohort 1 were recovered from the Virginia Mason Medical Center Central Processing Lab after all tests required for clinical care were complete under approval by the Benaroya Research Institute (BRI) protocol IRB20-036. This study was granted a waiver of informed consent as the study was considered minimal risk and due to the urgency of COVID-19 research at the time of sample collection. Samples for cohort 2 were collected from Harborview Medical Center under approval by University of Washington (UW) protocol UW IRB STUDY 09763. Subjects were eligible for inclusion if they were positive for SARS-CoV-2 infection by RT-PCR nasal swab test. Subjects in cohort 2 required admission to the ICU for enrollment. Number of hospital free days was calculated by subtracting the number of days hospitalized from 28. Individuals who were hospitalized for at least 28 days or died during the course of admission were categorized as 0. COVID severity score was based on the previously published definition (5, 17).

### Definition of sHLH in COVID-19

We adapted published criteria for HLH and MAS (18, 19) to create a definition of COVID-sHLH based on available clinical and laboratory data. Here, we defined COVID-sHLH as subjects with a ferritin > 1000 accompanied by cytopenias in 2 or more lineages (WBC < 5000 OR ANC < 1000, hemoglobin < 9 or hematocrit < 27, platelets < 100,000), and elevated transaminases (either AST or ALT > 30) OR subjects with a ferritin > 3000. Subjects for whom no ferritin was measured were classified in the no sHLH group (n= 31 for cohort 1, n=37 for cohort 2).

### Proteomics

#### Olink

For cohort 1, plasma was isolated from blood and stored at −80C. For safety protocols in place at Olink at the time of the study, samples were heat-inactivated and re-frozen prior to analysis. Samples were run on the following panels: CARDIOVASCULAR II (v.5006), CARDIOVASCULAR III (v.6112), IMMUNE RESPONSE (v.3203), IMMUNO-ONCOLOGY (v.3111), INFLAMMATION (v.3022). A preset list of targets of proteins involved in monocyte responses or hyperinflammatory responses was analyzed from the earliest available time point using a Mann Whitney U test. Those samples that did not pass internal QC at Olink were excluded from subsequent analysis.

#### SomaScan

For cohort 2, blood was collected in EDTA anticoagulated tubes within 24 hours of ICU admission. Plasma was isolated and aliquoted immediately. All samples underwent two freeze-thaw cycles prior to analysis. Plasma samples were analyzed with aptamer-based proteomic profiling using the SomaScan^®^ Platform which measures 4,984 human proteins with high specificity. The assays were performed as previously described (20). Concentrations of each protein bound to its corresponding aptamer was reported as relative fluorescent units (RFU). Median intra- and inter-assay coefficients of variation are approximately 5%. Two subjects had samples that failed quality control thresholds and were excluded from further analysis. After selection for participants positive for COVID-19, 101 remaining participants were included in analyses.

#### Meso Scale Discovery (MSD)

sPD-L1 protein concentration was measured in patient plasma collected in EDTA tubes at site 2 using an electrochemiluminescent immunoassay per the manufacturer’s protocol (Meso Scale Discovery) [R-Plex sPD-L1 (F214C)]. Plasma samples underwent two freeze-thaw cycles prior to analysis. All analytes measured met quality control parameters including: 1) intraplate % CV <20%; 2) interplate % CV <20%, and 3) <10% of samples with measurement below the lower limit of detection. Concentrations were calculated from a standard curve per manufacturer’s protocol.

### RNA sequencing

Whole blood was obtained from subjects in cohort 1 and PBMCs were isolated using Ficoll-Paque (VWR) per manufacturer’s instructions and frozen. Thawed PBMCs were stained with antibodies (Supplemental Table S1). Live, CD3-CD19-CD56-CD15-CD14+ x HLA-DR+ cells were sorted on a FACSAria Fusion (BD Biosciences) into media containing 10% human serum. RNA was isolated using RNeasy Micro kit (Qiagen). RNA libraries were built using SMART-Seq v4 Ultra Low Input RNA Kit for Sequencing (Takara) and analyzed by RNA sequencing as previously described (21). To detect differentially expressed genes between sorted cell subsets, the RNA-seq analysis functionality of the linear models for microarray data (Limma) R package was used. A false discovery rate adjustment was applied to correct for multiple testing.

### Genetics

5 μg of DNA extracted from whole blood of 8 deceased subjects from site 2 who met our definition for sHLH was submitted to Invitae for the Primary Immunodeficiency panel (test code 08100).

### Statistical analysis

Clinical data parameters were compared by the relevant statistical test (Mann Whitney U test, Welch’s t-test, or Chi-square) as indicated. O-link and MSD data were analyzed using Mann Whitney U test. SomaScan data was analyzed with linear regression of the log_2_ transformation of within subject mean normalized RFU values as the dependent variable and sHLH status as a covariate. This results in the beta coefficient of sHLH status being the log_2_ fold change between case and control groups. Analyses were covariate adjusted for age, sex, BMI. Type I error was mitigated using a false discovery rate (FDR<0.05 threshold derived using the Benjamini-Hochberg method).

## Results

### COVID-sHLH criteria and study populations

Similar proportions of subjects in both cohorts met our COVID-sHLH criteria as defined above. In cohort 1 (n=85), 7 subjects (8.2%) met the definition for COVID-sHLH (Table 1). In cohort 2 (n=101), 10 subjects (9.9%) met this definition. Subjects with and without sHLH in both cohorts were of similar mean age and BMI (Table 1). Subjects in cohort 1 with sHLH were more likely to be male (71.4%) as compared to those without sHLH (47.4%) but this difference was not seen in cohort 2 which was male predominant (Table 1). In both cohorts, history of autoimmunity, cancer, COPD, asthma, and organ transplantation were low. Both cohorts had higher rates of chronic kidney disease in subjects with COVID-sHLH compared to those without sHLH; cohort 1 (28.6% vs 10.3%) and cohort 2 (30% vs 16.5%). Rates of hypertension and diabetes were higher in subjects with COVID-sHLH from cohort 1 compared to those without sHLH (hypertension 57.1% vs 29.4%, diabetes 71.4% vs 46.1%), whereas in cohort 2 rates of hypertension and diabetes were similar regardless of whether subjects had sHLH or not (Table 1).

**Table 1:** Cohort Demographics.

|  | **Cohort 1** |  | **Cohort 2** |  |
| --- | --- | --- | --- | --- |
| sHLH Features | No | Yes | No | Yes |
| Number, % | 78 (91.8%) | 7 (8.2%) | 91 (90.1%) | 10 (9.9%) |
| Mean age in years (SD) | 59 (17.9) | 63 (6.5) | 54.5 (15.9) | 58.4 (17.9) |
| # Males (%) | 37 (47.4%) | 5 (71.4%) | 65 (71.4%) | 8 (80%) |
| **Race** |  |  |  |  |
| American Indian | 6 (7.7%) | 0 (0%) | 5 (5.5%) | 3 (30%) |
| Asian | 7 (9.0%) | 0 (0%) | 7 (7.7%) | 0 (0%) |
| Black | 9 (11.5%) | 3 (42.9%) | 9 (9.9%) | 0 (0%) |
| White | 36 (46.2%) | 1 (14.3%) | 60 (65.9%) | 7 (70%) |
| Unknown | 20 (25.6%) | 3 (42.9%) | 9 (9.9%) | 0 (0%) |
| **BMI** (mean, SD) | 30.1 (8.3) | 33.4 (9.3) | 29.9 (7.6) | 28.6 (7.5) |
| **Comorbidities** |  |  |  |  |
| Hypertension | 23 (29.4%) | 4 (57.1%) | 42 (46.1%) | 6 (60%) |
| Diabetes | 22 (28.2%) | 5 (71.4%) | 29 (31.9%) | 4 (40%) |
| Autoimmune disease | 7 (9.0%) | 1 (14.3%) | 5 (5.5%) | 2 (20%) |
| Cancer | 11 (14.1%) | 0 (0%) | 9 (9.9%) | 1 (10%) |
| Chronic kidney disease | 8 (10.3%) | 2 (28.6%) | 15 (16.5%) | 3 (30%) |
| Asthma | 8 (10.3%) | 1 (14.3%) | 12 (13.2%) | 0 (0%) |
| COPD | 9 (11.5%) | 0 (0%) | 8 (8.8%) | 0 (0%) |
| Organ transplant | 1 (1.3%) | 0 (0%) | 2 (2.2%) | 0 (0%) |
| **Clinical labs\*** |  |  |  |  |
| Min WBC | 6.5 (4.5-8.3) | 3.9 (2.6-4.45) | 6.2 (4.0-7.89) | 5.69 (4.71-9.21) |
| Min ANC | 4.4 (2.5-5.8) | 2.45 (2.06-3.25) | 5.06 (2.77-8.35) | 6.25 (3.69-10.11) |
| Min Hemoglobin | 11 (12.8-8.9) | 11.3 (11-11.8) | n/a | n/a |
| Min Hematocrit | n/a | n/a | 28 (24-32) | 24 (20.75-29.75) |
| Min Platelets | 177 (128.8-244) | 140 (94.5-156.3) | 164 (131.5-202) | 147.5 (79-212.2) |
| Max AST | 41 (27-61) | 128 (87-242) | 68 (37-94) | 91.5 (48-125) |
| Max ALT | 37 (21-65) | 78 (64-135) | 54 (34-99) | 32 (20-140) |
| Max Ferritin | 612 (235-1410)<sup>&</sup> | 3612 (3125-6835) | 650 (255-996)<sup>#</sup> | 4664 (1769-9209) |
| **Received Steroids** | 8 (10.3%) | 1 (14.3%) | 48 (52.7%) | 2 (20%) |
| **Received Tocilizumab** | 12 (15.4%) | 3 (42.9%) | 5 (5.5%) | 0 (0%) |

\* Median value (interquartile range), for cohort 2 median from within first 15 days of hospitalization
<sup>&</sup>n=47 for cohort 1
<sup>#</sup>n=54 for cohort 2

As expected based on the criteria for sHLH, ferritin was significantly higher in subjects with sHLH (median of 3612) compared to subjects without sHLH (median 612) in cohort 1 (p=0.016). Ferritin was numerically but not statistically higher in subjects with sHLH (median 4664) in cohort 2 compared to subjects without sHLH (median 649, p=0.15). Subjects with sHLH in cohort 1 were as likely to have received steroids (14.3%) as those without sHLH (10.3%) whereas subjects with sHLH in cohort 2 were less likely to have received steroids (20%) than those without COVID-sHLH (52.7%). Of note, clinical practice regarding the use of steroids in the treatment of COVID-19 changed over the course of the study period following publication of the RECOVERY trial (22).

Mortality was significantly higher in subjects with sHLH in cohort 2 compared to subjects without sHLH (70% vs 25.2%, p<0.01, Table 2). Mortality in subjects with sHLH tended to be higher in cohort 1 compared to subjects without sHLH (28.6 vs 11.5%, Table 2), although this difference was not statistically significantly different (p =0.2). Overall mortality was substantially higher in cohort 2 (29.7% mortality in cohort 2 compared to 12.9% mortality in cohort 1) reflective of the greater proportion of patients with high disease severity scores in cohort 2. Similarly, subjects in cohort 1 had more hospital free days than subjects in cohort 2 with the highest proportion of hospital free days amongst those without sHLH from cohort 1 (Table 2). Notably, subjects with sHLH in both cohorts had fewer hospital free days reflecting the higher severity of illness in these subjects (Table 2). Taken together, we present a definition of sHLH in COVID-19 infected patients and find these patients were more likely to have longer lengths of hospital stay and higher mortality.

**Table 2:**
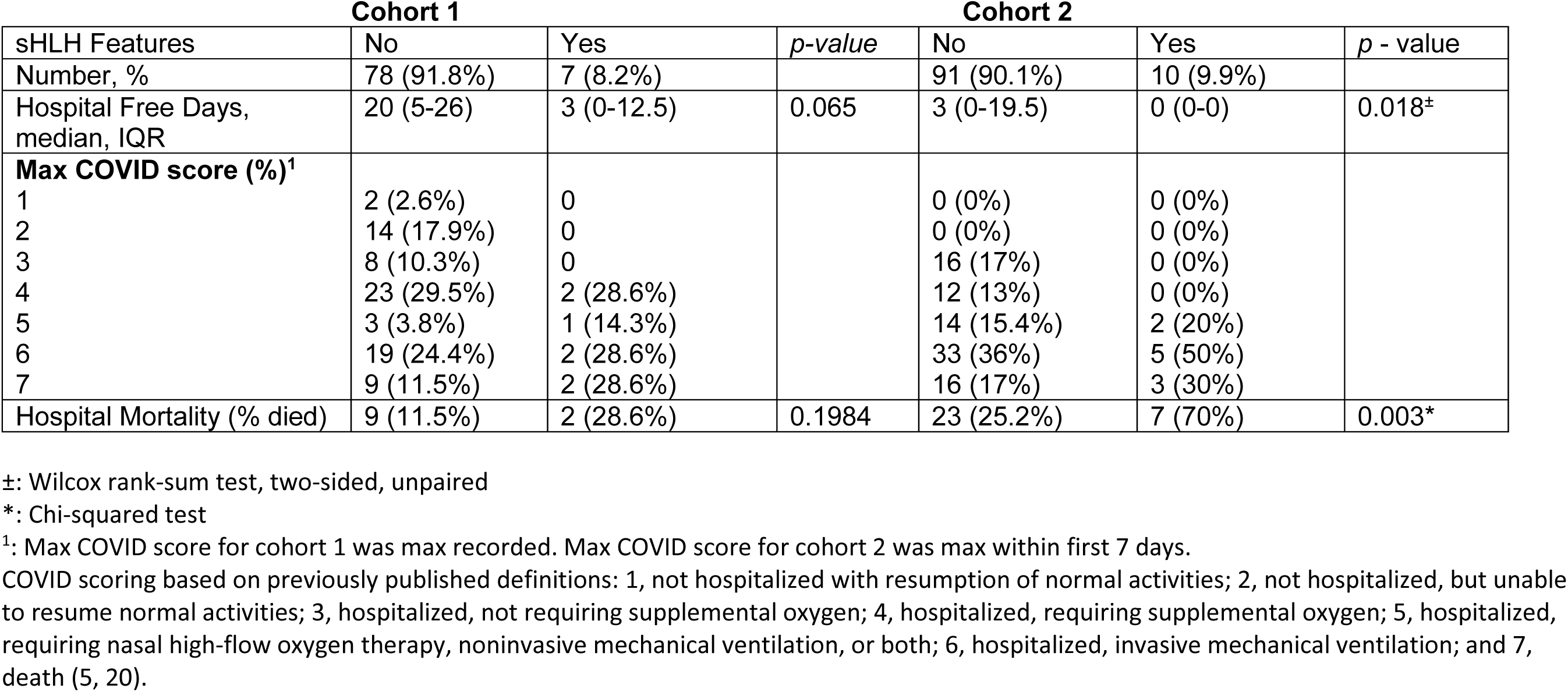
Cohort Outcomes.

| sHLH Features | Cohort 1 |  | *p-value* | Cohort 2 |  | *p - value* |
| --- | --- | --- | --- | --- | --- | --- |
|  | No | Yes |  | No | Yes |  |
| Number, % | 78 (91.8%) | 7 (8.2%) |  | 91 (90.1%) | 10 (9.9%) |  |
| Hospital Free Days, median, IQR | 20 (5-26) | 3 (0-12.5) | 0.065 | 3 (0-19.5) | 0 (0-0) | 0.018<sup>±</sup> |
| **Max COVID score (%)<sup>1</sup>** |  |  |  |  |  |  |
| 1 | 2 (2.6%) | 0 |  | 0 (0%) | 0 (0%) |  |
| 2 | 14 (17.9%) | 0 |  | 0 (0%) | 0 (0%) |  |
| 3 | 8 (10.3%) | 0 |  | 16 (17%) | 0 (0%) |  |
| 4 | 23 (29.5%) | 2 (28.6%) |  | 12 (13%) | 0 (0%) |  |
| 5 | 3 (3.8%) | 1 (14.3%) |  | 14 (15.4%) | 2 (20%) |  |
| 6 | 19 (24.4%) | 2 (28.6%) |  | 33 (36%) | 5 (50%) |  |
| 7 | 9 (11.5%) | 2 (28.6%) |  | 16 (17%) | 3 (30%) |  |
| Hospital Mortality (% died) | 9 (11.5%) | 2 (28.6%) | 0.1984 | 23 (25.2%) | 7 (70%) | 0.003* |

±: Wilcoxon rank-sum test, two-sided, unpaired
\*: Chi-squared test
<sup>1</sup>: Max COVID score for cohort 1 was max recorded. Max COVID score for cohort 2 was max within first 7 days.
COVID scoring based on previously published definitions: 1, not hospitalized with resumption of normal activities; 2, not hospitalized, but unable to resume normal activities; 3, hospitalized, not requiring supplemental oxygen; 4, hospitalized, requiring supplemental oxygen; 5, hospitalized, requiring nasal high-flow oxygen therapy, noninvasive mechanical ventilation, or both; 6, hospitalized, invasive mechanical ventilation; and 7, death (5, 20).

### sPD-L1, sTNF-R1 and IL-18BP as biomarkers of COVID-sHLH

We selected a set of candidate proteins to test for association with COVID-sHLH based on previous associations with sHLH/MAS in other conditions or based on activity in the IL-1/IL-18 or IFN pathways (23–28). These proteins were measured from plasma using the Olink platform for subjects in cohort 1. Levels of soluble CD163 (sCD163), IL-18 binding protein (IL-18BP), soluble tumor necrosis factor receptor 1 (sTNFR1), and sPD-L1 were significantly higher in subjects with COVID-sHLH compared to subjects without COVID-sHLH (p-value < 0.05, Fig. 1A). In contrast, we found no significant difference in levels of TNF (Fig. 1A). In subjects with sHLH compared to those without sHLH, the normalized protein expression (NPX) values were higher in subjects with sHLH compared to those without sHLH for sCD163 (9.10 vs 8.72, p=0.04), IL-18-BP (8.07 vs 7.35, p=0.01), sTNFR1 (9.46 vs 8.53, p=0.02), and sPD-L1 (8.44 vs 7.80, p=0.01).

**Figure 1.**
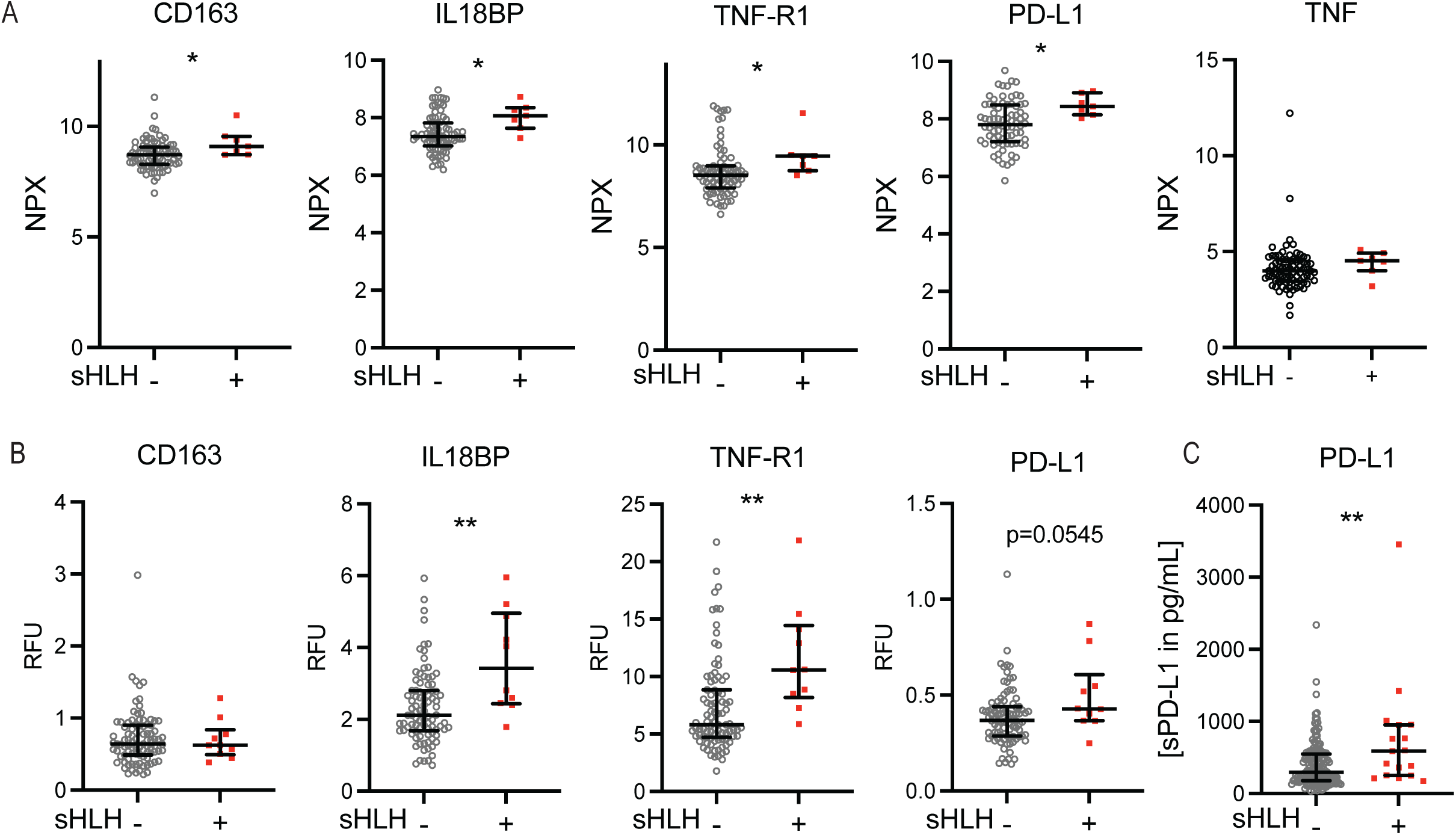
sPD-L1, sTNF-R1, and IL-18BP are elevated two independent cohorts of COVID-sHLH subjects. (A) Plasma samples (n=85) from cohort 1 were analyzed on the Olink platform and displayed as intensity normalized values. A.U. indicates arbitrary unit. Open circles: no sHLH (n=78, except for PD-L1 and TNF n=77), red squares: with sHLH (n=7). (B) Plasma samples from cohort 2 were analyzed using the SomaScan platform (n=101) and displayed as normalized RFU. Open circles: no sHLH (n=91); red squares: with sHLH (n=10). (C) Plasma samples from cohort 2 were analyzed for sPD-L1 (n=181) using an MSD assay. Open circles: no sHLH, red squares (n=164): with sHLH (n=17). Bars indicate median and interquartile range. * p < 0.05, ** p< 0.01 by two-tailed Mann-Whitney test.

To validate whether these proteins were associated with our defined sHLH criteria in an independent cohort, we analyzed levels of sCD163, IL-18BP, sTNFR1, and sPD-L1 in cohort 2 using the SomaScan platform. We found that there was a significantly higher level of sTNF-R1 and IL-18BP and a trend towards a significant difference in sPD-L1 (p=0.0545) in subjects with COVID-sHLH compared to subjects without sHLH (Fig. 1B). However, sCD163 levels were not significantly different between groups in cohort 2 (Fig. 1B). In order to further assess the association between sPD-L1 and COVID-sHLH, we measured this protein on the MSD platform which provides a specific protein concentration measurement, as opposed to arbitrary units, in a larger cohort of COVID-19 patients from site 2 (n=181). sPD-L1 was significantly higher in COVID-sHLH using MSD (Fig. 1C).

### COVID-sHLH protein and pathway discovery analysis

Given identification of several potential COVID-sHLH biomarkers using a targeted approach, we pursued an unbiased approach using the data from 4984 aptamers measured by SomaScan in cohort 2 to identify novel protein associations with sHLH. Multiple (n=185) proteins were significantly differentially expressed between subjects with and without sHLH (FDR<0.05) (Fig. 2A, Supplemental Table S2), with n=110 proteins higher in COVID-sHLH and n=75 proteins higher in those without sHLH. CHCHD10, coiled-coil-helix-coiled-coil-helix domain containing 10, was one of the most significantly differential expressed proteins and was increased in COVID-sHLH (Fig. 2A-B). CHCHD10 encodes a mitochondrial protein that may play a role in maintenance of cristae morphology or oxidative phosphorylation and has not been previously associated with sHLH. To validate this finding, we analyzed CD14+ monocytes isolated from subjects in cohort 1 (n=25) using RNA sequencing and found that CHCHD10 transcripts were significantly increased in COVID-sHLH monocytes (Fig. 2C, FDR = 0.017). Notably, this was the only significant differentially expressed gene in this small cohort of COVID-sHLH (n=3) compared to those without sHLH (n=22). UNC13B, a homolog of the HLH-associated gene UNC13D, trended towards differential expression between those with and without sHLH (Fig. 2C, FDR = 0.147).

**Figure 2.**
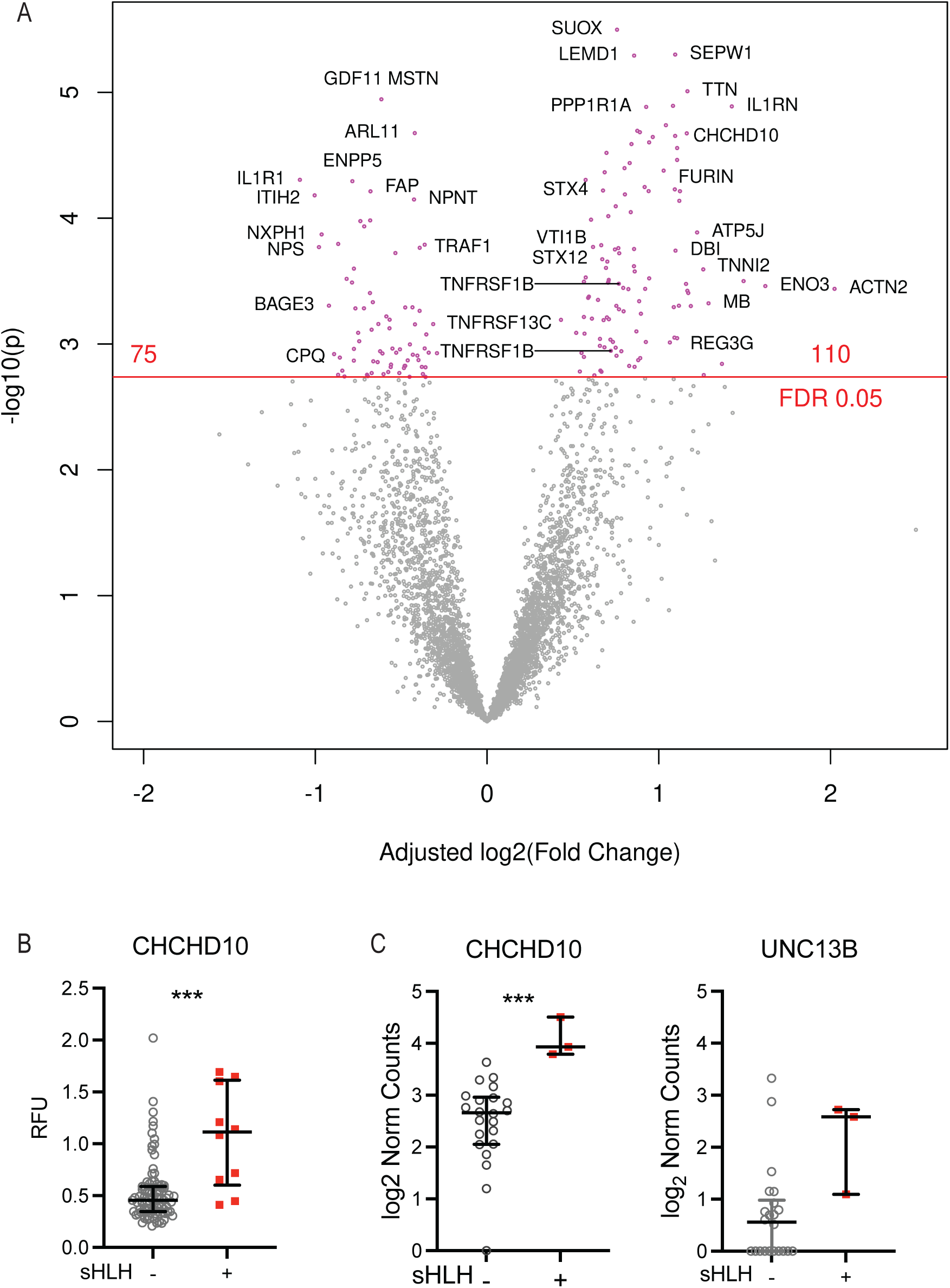
Syntaxin and IL-1 pathway proteins and CHCHD10 are differentially expressed in COVID-sHLH subjects. (A) Plasma samples from cohort 2 were analyzed using SomaScan platform (n=101; no sHLH n=91; sHLH n=10). Volcano plot displays differentially expressed proteins between subjects with sHLH vs those without sHLH adjusted for age, sex, and BMI. FDR < 0.05. (B) Plasma levels of CHCHD10 from cohort 2 displayed as normalized RFU from SomaScan data presented in Panel A. Bars indicate median and interquartile range. (C) Normalized read counts of transcripts from RNA sequencing of CD14+ monocytes from subjects without sHLH (n=22) or subjects with sHLH (n=3) from cohort 1. Open circles: no sHLH, red squares: with sHLH. Bars indicate median and interquartile range. *** p < 0.001 by two-tailed Mann-Whitney test.

We then used pathway analysis to discover relationships between serum proteins identified in the SomaScan analysis (Fig. 2A). Using STRING analysis, we found that proteins significantly higher in COVID-sHLH were not randomly associated with one another (protein-protein interaction enrichment p value < 1.0 x 10^-16^). Using gene ontology (GO) cellular component analysis, the SNARE complex, extracellular vesicle, and extracellular exosome terms were identified among proteins significantly enriched in subjects with COVID-sHLH (Table 3). Identification of the SNARE complex was intriguing given the known association of syntaxin genes with HLH (29–31). Syntaxin proteins (STX4, STX12, STX8) and the vesicle transport protein (VTI1B, vesicle transport through interaction with t-SNAREs 1B) were among the most significantly increased in samples from COVID-sHLH subjects (Fig. 2A). In addition to identification of the SNARE complex and syntaxin proteins involved in vesicle transport, GO pathway analysis identified IL-1 mediated signaling, positive regulation of protein secretion, and multiple vesicle fusion pathways as significant pathways enriched in COVID-sHLH (Supplemental Table S3).

**Table 3:**
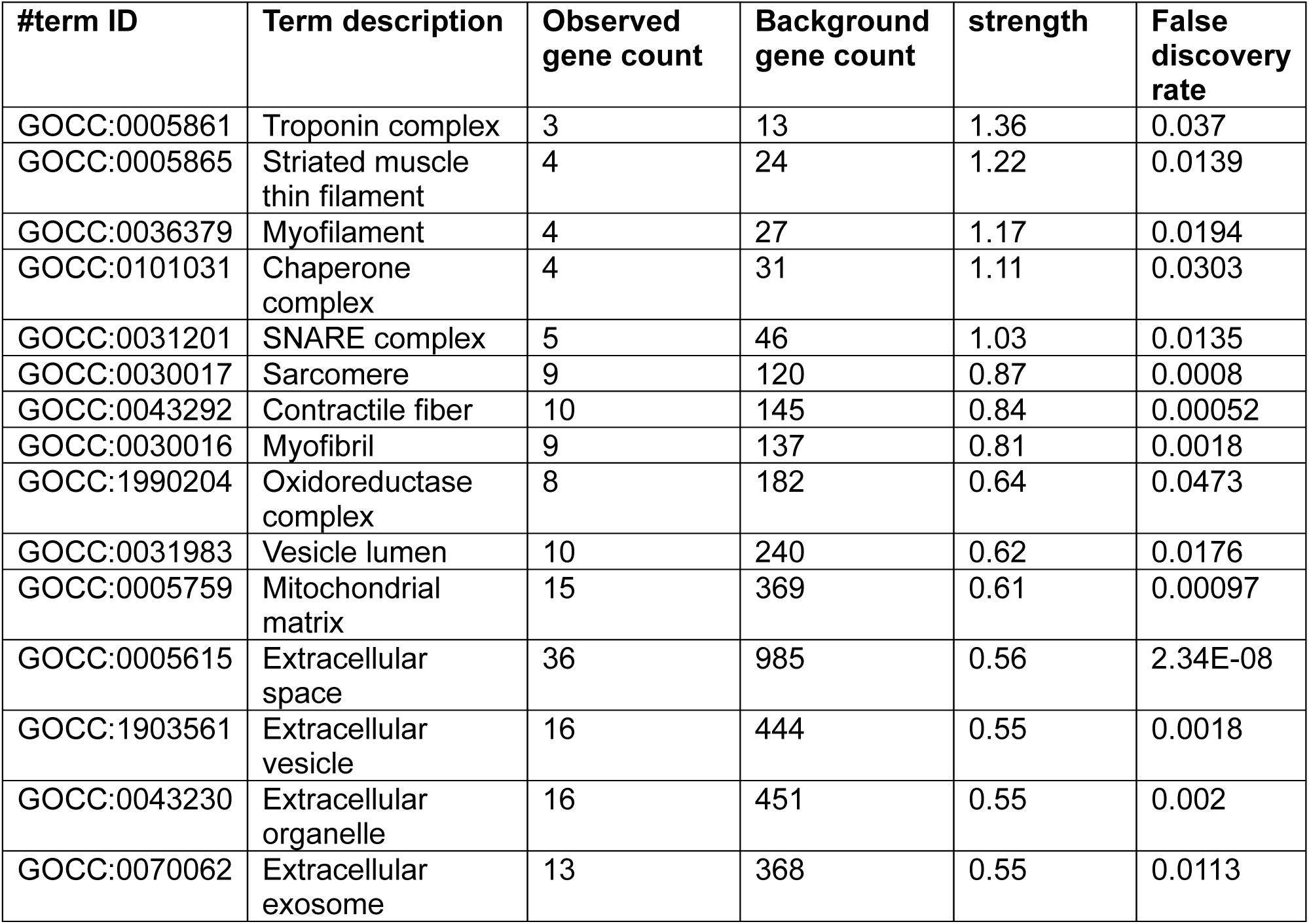
GO Cellular Component Analysis.

| #term ID | Term description | Observed gene count | Background gene count | strength | False discovery rate |
| --- | --- | --- | --- | --- | --- |
| GOCC:0005861 | Troponin complex | 3 | 13 | 1.36 | 0.037 |
| GOCC:0005865 | Striated muscle thin filament | 4 | 24 | 1.22 | 0.0139 |
| GOCC:0036379 | Myofilament | 4 | 27 | 1.17 | 0.0194 |
| GOCC:0101031 | Chaperone complex | 4 | 31 | 1.11 | 0.0303 |
| GOCC:0031201 | SNARE complex | 5 | 46 | 1.03 | 0.0135 |
| GOCC:0030017 | Sarcomere | 9 | 120 | 0.87 | 0.0008 |
| GOCC:0043292 | Contractile fiber | 10 | 145 | 0.84 | 0.00052 |
| GOCC:0030016 | Myofibril | 9 | 137 | 0.81 | 0.0018 |
| GOCC:1990204 | Oxidoreductase complex | 8 | 182 | 0.64 | 0.0473 |
| GOCC:0031983 | Vesicle lumen | 10 | 240 | 0.62 | 0.0176 |
| GOCC:0005759 | Mitochondrial matrix | 15 | 369 | 0.61 | 0.00097 |
| GOCC:0005615 | Extracellular space | 36 | 985 | 0.56 | 2.34E-08 |
| GOCC:1903561 | Extracellular vesicle | 16 | 444 | 0.55 | 0.0018 |
| GOCC:0043230 | Extracellular organelle | 16 | 451 | 0.55 | 0.002 |
| GOCC:0070062 | Extracellular exosome | 13 | 368 | 0.55 | 0.0113 |

In cohort 2, IL-1RN (also known as IL-1 receptor antagonist or IL-1RA), the endogenous inhibitor of IL-1, was significantly elevated in COVID-sHLH whereas IL-1R1 (also known as IL-1 receptor type 1) was significantly lower in COVID-sHLH by SomaScan (Fig. S1A). Using O-link available data from cohort 1, we did not replicate these findings with no significant difference in IL-1RN and an increase, rather than reduction, in IL-1R1 (Fig S1B). It is possible that this discrepancy may be due to differences in targets between platforms or differences in disease severity between the two cohorts. These findings provide mixed evidence for the importance of the regulation of the IL-1 pathway in COVID-sHLH that requires further exploration. Clinical studies of anakinra, recombinant IL-1RN, in COVID-19 have been mixed with a phase 2/3 RTC suggesting no improvement in outcomes with severe COVID pneumonia (32).

### Genetics of COVID-sHLH

There is increasing evidence that genetic variants in immune pathways may contribute to dysregulated responses to infectious triggers including COVID-19 (9, 33–36). In addition, we identified an association between the SNARE complex pathway and COVID-sHLH. Genetic variants in the SNARE complex, such as STX11 and STXBP2, have previously been identified as a cause of pHLH. We therefore hypothesized that COVID-sHLH subjects might have genetic variants in immune genes that placed them at increased risk of fatal outcomes. Deceased subjects from site 2 who met our definition of COVID-sHLH and who had sufficient DNA available for sequencing were analyzed using Invitae’s Primary Immunodeficiency Panel. Of the 8 subjects analyzed, 5 subjects (62.5%) had variants of unknown significance (VUS) or pathogenic mutations in genes known to be associated with primary immunodeficiencies, including two heterozygous pathogenic mutations, both in genes known to cause autosomal recessive disease, and 10 VUS (Supplemental Table S4, Table 4). Then, we excluded VUS predicted to be benign or likely benign, based on ACMG classification. Three subjects (37.5%) contained at least one variant that was predicted to be damaging (Table 4). Pathogenic heterozygous mutations in *TMPRSS15* and *DOCK8* were identified in these deceased COVID-sHLH subjects. Homozygous or compound heterozygous mutations in *TMPRSS15* lead to enterokinase deficiency which causes intestinal malabsorption, chronic diarrhea and failure to thrive. Homozygous mutations in *DOCK8* lead to autosomal recessive hyper-IgE syndrome with combined immunodeficiency which is associated with recurrent infections among other manifestations. VUS in genes that were predicted to be deleterious or probably damaging were identified in *MS4A1* and *TAOK2*. No variants were identified in genes known to be associated with HLH (e.g. *PRF1*, *STX11*, *STXBP2*, *UNC13D*, *SH2D1A*, *XIAP*, *LYST*, *AP3B1*). Taken together, these data suggest that variants in immune-related genes may affect susceptibility to COVID-sHLH and fatal outcomes, although we identified no known HLH variants.

**Table 4:**
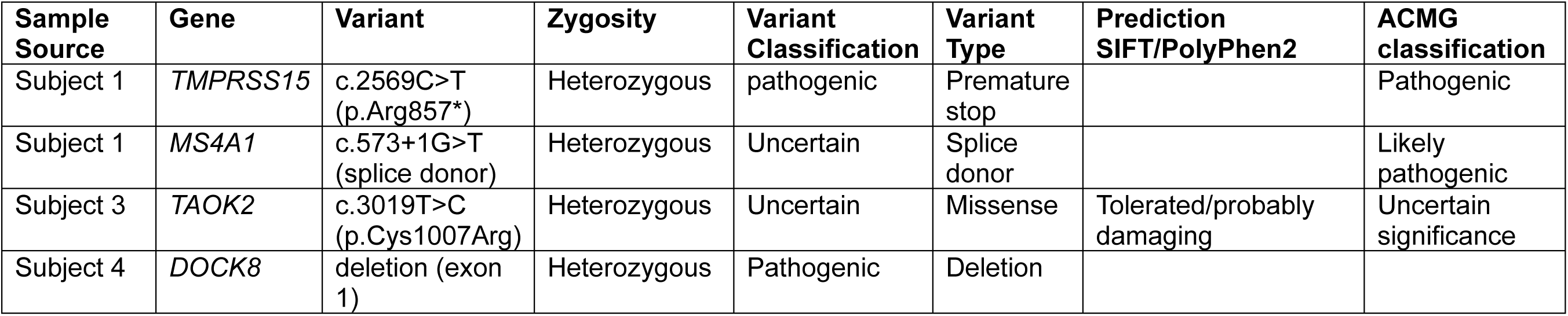
Mutations identified in deceased COVID-19 subjects with sHLH.

## Discussion

We utilized available clinical data to identify subjects with COVID-sHLH based on elements from consensus criteria to define novel pathways in COVID-19 associated sHLH. Our study further supports a role for proteins previously associated with other forms of sHLH including IL-18BP and sTNF-R1 and identifies novel biomarkers such as sPD-L1 and pathways of disease including the syntaxin pathways in COVID-sHLH. We also identified variants in several genes involved in immune responses, suggesting that genetic alterations in immune-related genes may contribute to hyperinflammation and fatal outcomes in COVID-19, although we did not identify any HLH-associated variants in individuals with fatal COVID-19.

Here we identify IL-18BP, sTNF-R1, and sPD-L1 as potential biomarkers for COVID-sHLH in two distinct cohorts using different platforms for protein analysis. Although the proteomic data from the two cohorts were generated using different techniques, the reproducibility of these findings across multiple platforms lends strength to these findings. IL-18BP, sTNF-R1, and sPD-L1 all act to dampen inflammatory responses and are induced by IFNs. sPD-L1 and IL-18BP are primarily induced by IFNγ (14, 37). IL-18BP binds the proinflammatory cytokine, IL-18, to reduce its activity by preventing receptor binding. sTNF-R1 can bind circulating TNF to inhibit its pro-inflammatory signaling. sPD-L1 is thought to be produced by proteolytic cleavage of the membrane bound protein PD-L1 but retains the ability to inhibit T cell activation and cytokine production. IL-18BP and sTNF-R1, but not sPD-L1, have been previously reported to be elevated in subjects with MAS/HLH (23, 24), whereas higher levels of sPD-L1 and sTNF-R1 are reported to be associated with severe disease in COVID-19 patients, need for mechanical ventilation, and fatal outcomes (7, 15, 16, 38, 39). To our knowledge, sPD-L1 has not been previously identified as a biomarker in HLH or MAS but given that it is induced by IFNγ and the critical role that IFNγ plays in HLH/MAS, sPD-L1 may prove to be a useful biomarker for other hyperinflammatory conditions. While elevated levels of sPD-L1, sTNF-R1, and IL-18BP have been noted in patients with severe COVID-19, none of these proteins have not been previously identified as biomarkers in COVID-sHLH.

In comparison to our work, a previous study has identified several serum proteins that may distinguish critical COVID-19 illness from non-COVID-19 sHLH/MAS (40). The authors found IFNγ was the most effective marker at separating sHLH/MAS from critical COVID-19 (40). Notably, in this study, the subjects with COVID-19 were defined by disease severity rather than by HLH/MAS-like criteria (40), which may contribute to between our studies. We found significant increase in sPD-L1 levels, an IFNγ induced protein, in COVID-sHLH subjects from two independent cohorts, including in cohort 2 where all subjects were critically ill, supporting a role for the IFNγ pathway in COVID-sHLH.

Our data support a role for the syntaxin pathways in COVID-sHLH. Syntaxins are proteins that are involved in exocytosis and play important roles in membrane fusion to promote protein trafficking intracellularly and for release of vesicle contents extracellularly. Our data implicate the secretory pathway, vesicle trafficking and cytotoxic cell degranulation as important in COVID-sHLH similar to the critical role they play in HLH/MAS (41). Previous studies have demonstrated interactions between various syntaxin proteins and SARS-CoV-2 ORFs (42, 43) in order to promote viral infection of and trafficking in host cells. STX8, which was upregulated in COVID-sHLH subjects (Figure 2A), has been identified as a biomarker in deceased COVID subjects (44), further implicating components of the SNARE complex in severe COVID and corroborating our findings. Taken together, our data confirms the importance of the syntaxin pathway in SARS-CoV-2 infection and further highlights a significant role for this pathway in subjects with COVID-sHLH.

Although our data identify a role in the syntaxin pathway in patients with COVID-sHLH, we did not detect any variants in known HLH/MAS-associated genes in fatal COVID-sHLH subjects. Variants identified as pathogenic heterozygous mutation or that were predicted to be likely damaging in our COVID-sHLH cohort were detected in *TMPRSS15*, *DOCK8*, *MS4A1*, and *TAOK2*. Notably, here, we identify deletion of exon 1 in *DOCK8* in a subject with fatal COVID-19 and COVID-sHLH. Whether this deletion in *DOCK8* alters NK cell function is worthy of further investigation especially in light of recent data suggesting that *DOCK8* variants may alter NK cell degranulation (45). *DOCK8* variants have been proposed to be involved in some cases of MAS/HLH and cases of multisystem inflammatory syndrome in children (MIS-C), a post infectious hyperinflammatory syndrome in children recently infected with SARS-CoV-2. *DOCK8* variants identified in MIS-C were introduced into human NK cells by lentiviral infection and decreased NK cell degranulation (45). Together, these data suggest that *DOCK8* variants or mutations may affect NK cell function leading to hyperinflammatory syndromes. Further experiments are needed to determine whether the heterozygous deletion in exon 1 of *DOCK8* from this subject affects NK cell function.

We also identified a pathogenic heterozygous mutation in *TMPRSS15* in a deceased COVID-sHLH subject. Transmembrane serine protease 15 (TMPRSS15) is an enteropeptidase whose expression is restricted to the small intestine that has been reported to be increased in fecal samples with high levels of SARS-CoV-2 RNA (46). Intestinal expression of *TMPRSS15* may suggest a role for entry of SARS-CoV-2 virus in this tissue. Whether this variant could play a role in susceptibility to COVID-sHLH is unclear. Although we did not identify any variants in HLH-associated genes in this small cohort of fatal COVID-sHLH patients, other groups have identified such variants in severe COVID-19 patients. For example, variants in *PRF* (47), *UNC13D* (48), and *AP3B1* (48) and a monoallelic missense mutation in *STXBP2* (49) have been identified by other groups in patients with severe COVID-19.

The strengths of our study include the inclusion of two distinct patient cohorts from two different institutions and reproducible identification of protein biomarkers across distinct platforms. The limitations of our study include the small number of COVID-sHLH subjects, particularly those with DNA available for genetic analyses, and the geographic limitation of our study to a single US city with data collected during the first COVID wave. We were unable to define sHLH using existing consensus criteria or recently developed definitions for COVID-19 cytokine storm (1, 4) as many of the data included in these criteria were not available for our subjects. In spite of these limitations, we have identified novel biomarkers for COVID-sHLH, namely sPD-L1, sTNF-R1, and IL-18BP, all of which are IFN inducible proteins, highlighting the importance of IFNs in COVID-sHLH. Further, we have identified a potentially damaging mutation in *DOCK8* in a patient with fatal COVID-19, which, if functionally relevant for NK cell degranulation, would further highlight the potential of HLH/MAS/CSS pathophysiology in driving a subset of severe SARS-CoV-2 infections.

## Acknowledgements

The authors thank the study participants and their families and clinical teams caring for these patients during the COVID-19 pandemic. The authors gratefully acknowledge the expertise of the Benaroya Research Institute’s Clinical and Translational research groups, including Thien-Son Nguyen and the Center for Interventional Immunology, the Benaroya Research Institute COVID-19 Research Team for sample collection and processing in the early pandemic, Colin O’Rourke, Alyssa Ylescupidez and Tee Bahnson for clinical data abstraction and statistical support, Adam Wojno, PhD of the Cell and Tissue Analysis Group, Vivian Gersuk, PhD, Quynh-Anh Nguyen, and Kimberly O’Brien of the Genomics Core, and members of the Hamerman and Mikacenic labs. COVID-19 sample collection at Virginia Mason Medical Center was supported by the Benaroya Family Foundation, the Leonard and Norma Klorfine Foundation, Glenn and Mary Lynn Mounger, and Monolithic Power Systems. Olink data collection was supported by Biogen, Inc, who had no role in the analysis or preparation of this manuscript. We also thank the M.J. Murdock Charitable Trust for supporting equipment at the Benaroya Research Institute that enabled this work.

Funding for this study was provided by: NIH grant R01AI150178-01S1 (to JAH), U19AI142733 (to CM), R01HL149676 (to CM), F32 HL156516 (to SPC), NIH NHLBI T32 HL007287-42 (to SEH). SPC is also supported by the Arthritis National Research Foundation and a CARRA/Arthritis Foundation Career Development Award. The authors wish to acknowledge CARRA and the ongoing Arthritis Foundation financial support of CARRA.

## Supplemental Figure

**Figure S1: IL-1 pathway associated proteins are variably expressed between cohorts 1 and 2.**

(A) Plasma samples (n=101; no sHLH n=91; sHLH n=10) from cohort 2 were analyzed using SomaScan platform and displayed as normalized RFU. (B) Plasma samples (n=85; no sHLH n=77 for IL-1RN, n=78 for IL-1R1; sHLH n=7) from cohort 1 were analyzed using the Olink platform and displayed as intensity normalized values. Open circles: no sHLH, red squares: with sHLH. Bars indicate median and interquartile range. * p < 0.05 and *** p < 0.001 by two-tailed Mann-Whitney test.

