## Supplemental Data for "Identification of biomarkers for COVID-19 associated secondary hemophagocytic lymphohistiocytosis"

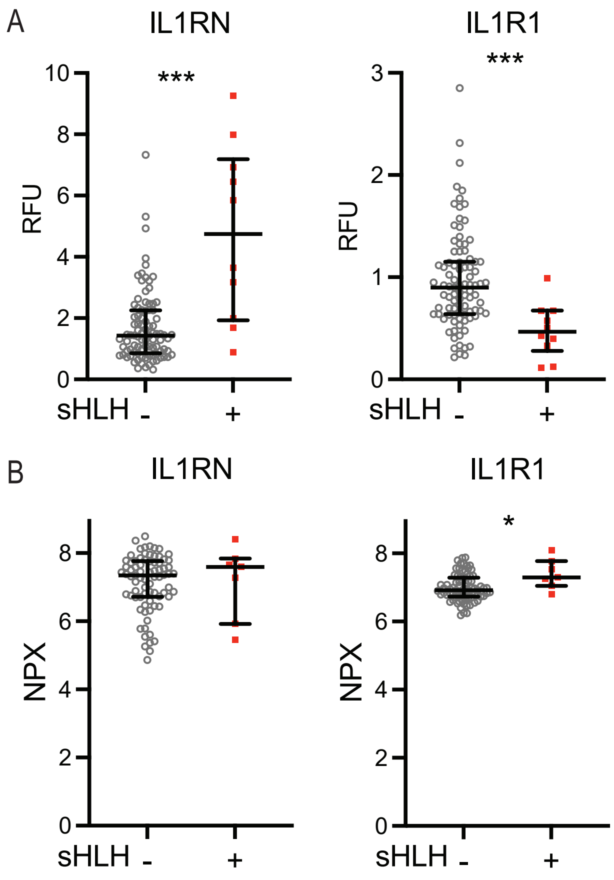

A
IL1RN
***
10
8
6
4
2
0
RFU
sHLH - +
IL1R1
***
3
2
1
0
RFU
sHLH - +
B
IL1RN
8
6
4
2
0
NPX
sHLH - +
IL1R1
*
8
6
4
2
0
NPX
sHLH - +


Figure S1

**Supplemental Table S1: Antibodies Used for Sorting Monocytes from PBMCs**

| Target | Fluorophore | Clone | Company |
|---|---|---|---|
| CD14 | BV510 | M5E2 | Biolegend |
| CD16 | PE | 3G8 | Biolegend |
| CD3 | BV650 | OKT3 | Biolegend |
| CD19 | BV650 | HIB19 | Biolegend |
| CD56 | BV650 | 5.1H11 | Biolegend |
| CD15 | BV650 | W6D3 | Biolegend |
| HLA-DR | APC | L243 | Biolegend |
| Live/dead | APC-E780 | | eBiosciences |

**Supplemental Table S3: GO Pathway Analysis**

| Pathway description | Protein aptamer count | False discovery rate |
|---|---|---|
| Striated muscle hypertrophy | 3 | 9.75E-04 |
| Cardiac myofibril assembly | 4 | 9.75E-04 |
| Muscle hypertrophy | 5 | 9.75E-04 |
| Negative regulation of heterotypic cell-cell adhesion | 5 | 9.75E-04 |
| IL-1 mediated signaling pathway | 50 | 0.001122 |
| Positive regulation of protein secretion | 49 | 0.001316 |
| Vesicle fusion with ER Golgi intermediate compartment (ERGIC) membrane | 3 | 0.00146 |
| Cornified envelope assembly | 4 | 0.001556 |
| Regulation of protein transport | 157 | 0.001836 |
| Synaptic vesicle fusion to presynaptic active zone membrane | 7 | 0.001922 |
| Vesicle fusion to plasma membrane | 7 | 0.001922 |

**Supplemental Table S4: Variants identified in deceased COVID-19 subjects with sHLH**

| Sample Source | Gene | Variant | Zygosity | Variant Classification | Variant Type | Prediction SIFT/PolyPhen2 | ACMG classification |
|---|---|---|---|---|---|---|---|
| Subject 1 | *TMPRSS15* | c.2569C>T (p.Arg857*) | Heterozygous | pathogenic | Premature stop | | Pathogenic |
| Subject 1 | *CARD8* | c.827A>T (p.His276Leu) | Heterozygous | Uncertain | Missense | Tolerated/benign | Likely benign |
| Subject 1 | *MS4A1* | c.573+1G>T (splice donor) | Heterozygous | Uncertain | Splice donor | | Likely pathogenic |
| Subject 1 | *TNFRSF11A* | c.29C>T (p.Pro10Leu) | Heterozygous | Uncertain | Missense | Tolerated/not available | Likely benign |
| Subject 2 | *TMEM173* | c.1124G>A (p.Arg375His) | Heterozygous | Uncertain | Missense | Deleterious/probably damaging | Likely benign |
| Subject 2 | *TNFRSF6B* | c.121G>T (p.Ala41Ser) | Heterozygous | Uncertain | Missense | Tolerated/benign | Likely benign |
| Subject 3 | *TAOK2* | c.3019T>C (p.Cys1007Arg) | Heterozygous | Uncertain | Missense | Tolerated/probably damaging | Uncertain significance |
| Subject 3 | *TNFRSF11A* | c.1805A>G (p.Glu602Gly) | Heterozygous | Uncertain | Missense | Tolerated/benign | Likely benign |
| Subject 4 | *DOCK8* | deletion (exon 1) | Heterozygous | Pathogenic | Deletion | | |
| Subject 4 | *STAT3* | c.645+13G>T (intronic) | Heterozygous | Uncertain | Uncertain | May disrupt consensus splice site | Benign |
| Subject 5 | *HYOU1* | c.1785G>A (silent) | Heterozygous | Uncertain | Silent | | Likely benign |
| Subject 5 | *NOD2* | c.392C>A (p.Ala131Asp) | Heterozygous | uncertain | Missense | | Likely benign |
